# Criticality of neuronal avalanches in human sleep and their relationship with sleep macro- and micro-architecture

**DOI:** 10.1101/2022.07.12.499725

**Authors:** Silvia Scarpetta, Niccolò Morrisi, Carlotta Mutti, Nicoletta Azzi, Irene Trippi, Rosario Ciliento, Ilenia Apicella, Giovanni Messuti, Marianna Angiolelli, Fabrizio Lombardi, Liborio Parrino, Anna Elisabetta Vaudano

**Affiliations:** Department of Physics “E.R.Caianiello”, University of Salerno, I-84084 Salerno, Italy; INFN, sez. di Napoli, gr. coll. Salerno, Italy; Nephrology, Dialysis and transplant unit, University Hospital of Modena, I-41121 Modena, Italy; Sleep Disorders Center, Department of Medicine and Surgery, University of Parma, I-43121 Parma, Italy; Department of Neurology, University of Wisconsin, Madison, WI-53705, USA; Department of Physics, University of Naples “Federico II”, I-80126 Napoli, Italy; Institute of Science and Technology Austria, A-3400 Klosterneuburg, Austria; Neurology Unit, Azienda Ospedaliero-Universitaria of Modena, OCB Hospital, Modena, Italy; Department of Biomedical, Metabolic and Neural Sciences, University of Modena and Reggio Emilia, I-4125 Modena, Italy

## Abstract

Sleep plays a key role in preserving brain function, keeping the brain network in a state that ensures optimal computational capabilities. Empirical evidence indicates that such a state is consistent with criticality, where scale-free neuronal avalanches emerge. However, the relationship between sleep, emergent avalanches, and criticality remains poorly understood. Here we fully characterize the critical behavior of avalanches during sleep, and study their relationship with the sleep macro- and micro-architecture, in particular the cyclic alternating pattern (CAP). We show that avalanche size and duration distributions exhibit robust power laws with exponents approximately equal to −3/2 e −2, respectively. Importantly, we find that sizes scale as a power law of the durations, and that all critical exponents for neuronal avalanches obey robust scaling relations, which are consistent with the mean-field directed percolation universality class. Our analysis demonstrates that avalanche dynamics depends on the position within the NREM-REM cycles, with the avalanche density increasing in the descending phases and decreasing in the ascending phases of sleep cycles. Moreover, we show that, within NREM sleep, avalanche occurrence correlates with CAP activation phases, particularly A1, which are the expression of slow wave sleep propensity and have been proposed to be beneficial for cognitive processes. The results suggest that neuronal avalanches, and thus tuning to criticality, actively contribute to sleep development and play a role in preserving network function. Such findings, alongside characterization of the universality class for avalanches, open new avenues to the investigation of functional role of criticality during sleep with potential clinical application.

**Significance statement:** We fully characterize the critical behavior of neuronal avalanches during sleep, and show that avalanches follow precise scaling laws that are consistent with the mean-field directed percolation universality class. The analysis provides first evidence of a functional relationship between avalanche occurrence, slow-wave sleep dynamics, sleep stage transitions and occurrence of CAP phase A during NREM sleep. Because CAP is considered one of the major guardians of NREM sleep that allows the brain to dynamically react to external perturbation and contributes to the cognitive consolidation processes occurring in sleep, our observations suggest that neuronal avalanches at criticality are associated with flexible response to external inputs and to cognitive processes, a key assumption of the critical brain hypothesis.

## 1 Introduction

Sleep is an active and dynamic complex process regulated by mechanisms that guide the alternation of non-Rapid Eye Movement (NREM) and REM sleep across the night. Physiologically, sleep macro-architecture is characterized by the concentration of deep slow wave sleep (SWS) (stage N3) in the first half of the night, and the dominance of light sleep (mainly N2) and REM sleep in the second half of the night, a balanced skewness modulated by the homeostatic process and by the REM-off and REM-on systems (Brown et al., 2012). Throughout the night numerous transitions among these sleep stages occur, and, within sleep stages, micro-states on the scale of seconds and minutes are observed.

The cyclic alternating pattern (CAP) is one of the major adaptive components of NREM sleep. According to Terzano et al (Terzano et al., 2000), CAP is a periodic EEG activity of NREM sleep characterized by repeated spontaneous phases of EEG activation (A phase) and subsequent phases of return to background activity (B phase), evolving in a cycling pattern. Based on the distribution of slow and fast EEG frequencies, the A phases of CAP are classified in three subtypes: A1, A2 and A3 (Terzano et al., 2002). These CAP subtypes are not randomly distributed along the night, but instead their appearance is linked with the homeostatic, ultradian and circadian mechanisms of sleep regulation (Parrino et al., 1993; Terzano et al., 2005). In particular, subtypes A1 are the expression of slow wave sleep propensity and follow the exponential decline of the homeostatic process. A covariance between A1 subtypes and sleep slow wave activity (SWA) has been proposed and the two electroencephalographic elements likely share the beneficial effect on sleep-related cognitive processes (Ferri et al., 2008; Aricò et al., 2010). Furthermore, both CAP-A1 subtype and SWA are involved in the build-up and maintenance of deep NREM sleep, acting as protectors for sleep continuity (Terzano et al., 2000; Parrino and Vaudano, 2018).

Spontaneous alternation of transient, synchronized active and quiescent periods is typical of systems that self-organize near a critical point of a non-equilibrium phase transition (Scarpetta and de Candia, 2014; Munoz, 2018; Lombardi et al., 2020b). Following a number of theoretical and numerical results (Cragg and Temperley, 1954; Crutchfield and Karl, 1990; Bak, 1996; Kinouchi and Copelli, 2006), it has been hypothesized that the brain self-organizes to criticality to maximize information processing and computational capabilities, and thus achieve optimal functional performance. This hypothesis is supported by empirical observations of neuronal avalanches — cascades of neural activity exhibiting power-law size and duration distributions——and long-range spatio-temporal correlations in neural activity across species, systems, and spatial scales (Linkenkaer-Hansen et al., 2001; Beggs and Plenz, 2003; Pasquale et al., 2008; Mazzoni et al., 2007; Petermann et al., 2009; Tagliazucchi et al., 2012; Palva et al., 2013; Ponce-Alvarez et al., 2018; Tkačik et al., 2015; Lombardi et al., 2021b; Mariani et al., 2021). In particular, presence of power law distributions indicates absence of characteristic temporal and spatial scales in the underlying dynamics, as observed at criticality.

Empirical evidence shows that neuronal avalanches during sleep exhibit power law size and duration distributions (Priesemann et al., 2013; Bocaccio et al., 2019; Allegrini et al., 2015), and that sleep may play an active role in tuning the brain to criticality (Meisel et al., 2017, 2013). At the same time, recent studies demonstrated that bursts of dominant cortical rhythms exhibit the hallmarks of self-organized critical dynamics across the sleep-wake cycle, suggesting that criticality could be essential mechanism for spontaneous sleep-stage and arousals transitions (Wang et al., 2019; Lombardi et al., 2020a). However, both the nature of the alleged criticality during sleep and the relationship between related avalanche dynamics and complex sleep macro- and micro-architecture—in particular the CAP—remain poorly understood. On the one hand, the scaling relations among exponents that are expected to hold at criticality have not been verified, and a general framework to understand criticality during sleep is currently missing. On the other hand, the dynamics of avalanches in connection with the highly variable and distinct states composing long- and short-term sleep cycles has not been studied, and the potential functional role of avalanches in sleep regulation has not been explored.

Herein, we fully characterize the critical behavior of neuronal avalanches during sleep, and determine the scaling relations that connect their critical exponents, showing that they are consistent with a specific universality class. We then study how avalanche dynamics interacts with the ascending and descending slope of the NREM-REM sleep cycles, and within NREM sleep, how the CAP phases couple with avalanche occurrence. Our analysis shows that avalanche dynamics is closely linked to NREM-REM sleep cycles across night sleep, and that neuronal avalanche occurrence correlates with the activation phase of the CAP. The results indicate that avalanches play an active role in sleep development, and point to a peculiar relationship between CAP, brain tuning to criticality during sleep, and cognitive processes.

## 2 Materials and Methods

### 2.1 Participants

The data analyzed in this study were extracted from overnight polysomnographic (PSG) recordings acquired from the Parma (Italy) Sleep Disorders Center database. Ten healthy subjects, 5 males and 5 females, mean aged 39,6 years (age range 28-53), were selected after the accomplishment of an entrance investigation. Subjects were selected based on the following inclusion criteria: (i) absence of any psychiatric, medical and neurological disorder (ii) normal sleep/wake habits without any difficulties in falling or remaining asleep at night: a personal interview integrated by a structured questionnaire confirmed good daytime vigilance level; (iii) no drug intake at the time of PSG and the month before; (iv) full night unattended PSG recordings performed with EOG (2 channels), EEG [Ag/AgCl electrodes placed according to the 10 - 20 International System referred to linked-ear lobes]. Recording electrodes were 19 (Fp2, F4, C4, P4, O2, F8, T4, T6, Fz, Cz, Pz, Fp1, F3, C3, P3, O1, F7, T3, T5) in seven subjects and 25 in the remaining three: (CP3, CP4, C5, C6, C2, C1, FC4, FC3, F4, C4, P4, O2, F8, T4, T6, Fz, Cz, Pz, F3, C3, P3, O1, F7, T3, T5), EMG of the submentalis muscle, ECG, and signal for SpO2 (pulse-oximetry O2-saturation). PSG recordings were acquired using a Brain Quick Micromed System 98 (Micromed, SPA). A calibration of 50 *μV* was used for EEG channels with a time constant of 0.1 s and a low-pass filter with 30 Hz cut-off frequency. EEG sampling rate was 256 Hz for six subjects while for the remaining four cases, one was recorded using a sampling rate of 128 Hz (subject #1) and the remaining three (subject #2, #3, #4) using 512 Hz. Each signal was recorded and examined by an expert clinician (CM, IT, LP). Analysis of sleep recordings (see Section 2.2) was performed with Embla RemLogic Software. The institutional Ethical Committee Area Vasta Emilia Nord approved the study (protocol nr. 19750).

### 2.2 Sleep analysis

#### Analysis of sleep macro-architecture

Sleep was scored visually in 30-s epochs using standard rules according to the American Academy of Sleep Medicine (AASM) criteria (Berry et al., 2017). Conventional PSG parameters included total time in bed (TIB) (minutes), total sleep time (TST) (minutes), sleep latency (minutes), rapid eye movement (REM) latency (minutes), sleep efficiency (%), wake after sleep onset (WASO) (minutes), as well as percentage of NREM (N1, N2, N3) and REM stages.

#### Analysis of sleep micro-architecture

Sleep micro-architecture evaluation refers to the quantification of CAP parameters based on the published international atlas (Terzano et al., 2002), and was manually performed using Embla REM-logic software by somnologists with strong expertise in the field (LP, CM). CAP is a global EEG phenomenon involving extensive cortical areas, thus CAP phases should be visible on all or most EEG leads. CAP is characterized by the alternation of phase A (transient electrocortical events) and phase B (low voltage background), both lasting between 2 and 60 seconds. According to published criteria (Terzano et al., 2002) phase A activities were classified into three subtypes:

1. **Subtype A1.** EEG synchrony is the predominant activity and the EEG desynchrony occupies < 20% of the whole phase A. Subtype A1 may include delta burst, K-complex sequences, vertex sharp transients, polyphasic bursts with < 20% of EEG desynchrony.
2. **Subtype A2.** It is a mixture of fast and slow rhythms where the EEG desynchrony occupies 20 – 50% of the entire phase A. This subtype includes polyphasic bursts with 20 – 50% of EEG desynchrony.
3. **Subtype A3.** EEG desynchrony is the predominant activity (> 50%) of the phase A. Subtype A3 includes K-alpha, EEG arousal and polyphasic bursts with > 50% of EEG desynchrony.

The percentage of NREM sleep occupied by CAP sequences defines the CAP rate. The absence of CAP for more than 60 seconds is scored as non-CAP, and represents the portion of NREM sleep characterized by a sustained physiologic stability. CAP sequences usually precede sleep stage transitions, and, specifically, subtypes A2 and A3 typically assist the shift from NREM to REM sleep. Under physiologic circumstances, CAP is not present during REM sleep. The following CAP variables were measured:

a. Total CAP time in minutes (total CAP time in NREM sleep),
b. CAP rate (the ratio of CAP time over total NREM sleep time),
c. Number and duration of CAP cycles,
d. Number and duration of each phase A subtype (A1, A2, A3),
e. Total number of phase A (derived by the sum of A1, A2, and A3),
f. Duration of phase A and B in seconds.

### 2.3 Neuronal avalanche analysis

Before performing avalanche analysis, waking and motion artifact segments during nocturnal sleep were manually identified and removed. Artifact-free EEG signals were z-score normalized to have zero mean and unit standard deviation (SD). To capture the spatio-temporal organization in avalanches of transient EEG events during sleep, we investigated clusters of large deflections of the artifact-free EEG signals. For each EEG channel, large positive or negative excursions beyond a threshold *θ* =±nSD were identified.

To define the threshold *θ*, we analyzed the distribution of EEG amplitudes (Fig. 1B). A Gaussian distribution of amplitudes is expected to arise from a superposition of many uncorrelated sources. Conversely, EEG amplitude distributions deviate from a Gaussian shape, indicating presence of spatio-temporal correlations and collective behaviors involving different cortical areas (Fig. 1). The comparison of the signal distribution to the best Gaussian fit indicates that the two distributions start to deviate from one another around *θ* = ±2 SD (Fig. 1). Thus, thresholds smaller than 2 SD would lead to the detection of many events related to noise in addition to real events whereas much larger thresholds will miss many of the real events. To avoid noise-related events while preserving most of relevant events, in this study we used a threshold value *θ* = ±2 SD. Importantly, avalanche distributions are robust for a wide range of threshold values > 2 SD (Supplementary Material, Fig. S1).

**Figure 1:**
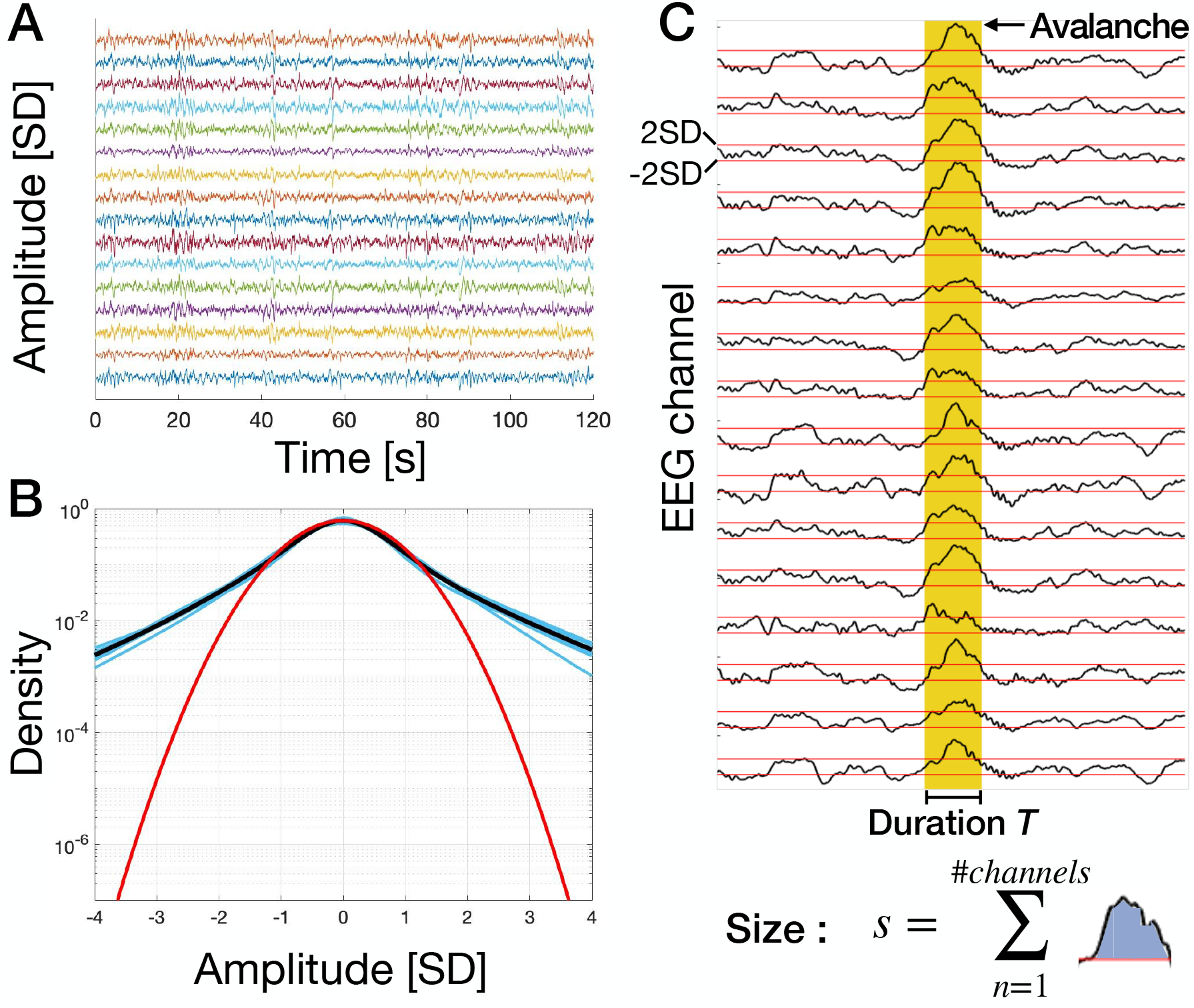
Identification of neuronal avalanches and definition of avalanche size and duration. (A) Segments (2 hours) of Z-score normalized EEG signal traces for an individual subject. Each trace correspond to an EEG channel. (B) Probability density of the z-score normalized EEG signal amplitude. The cyan curves in the background are the probability densities for all individual subjects (n = 10 subjects; for each subject we pooled all individual EEG channels). The black curve is the grand average over all subjects. The red curve is the best Gaussian fit for the grand average. We notice that the empirical probability density starts deviating from the Gaussian fit around ±2 SD. (C) A neuronal avalanche is defined as a continuous sequence of signal excursions beyond threshold (red thick line) on one or more EEG channels (upper panel). An avalanche is preceded and followed by periods in which EEG signal are below the threshold in all channels. The size of an avalanche is defined as the sum over all channels of the absolute values of the signals exceeding the threshold (bottom panel).

**Figure 2:**
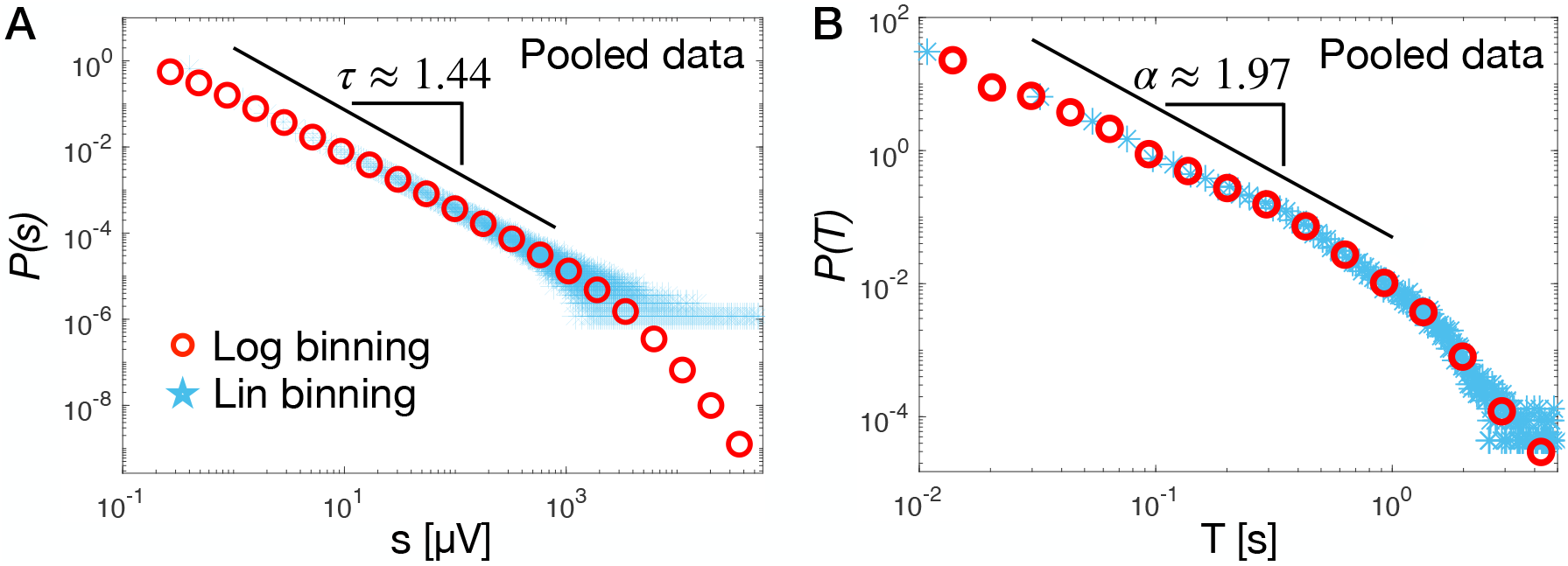
Avalanche size and duration distributions exhibit a robust power law behavior during sleep periods. (A) The distribution of avalanche sizes (red circles) follows a power law with exponent *τ* = 1.438 ±0.001 (fit ± std. error on the fit; pooled data, 10 subjects). The power law regime is followed by an exponential cut off. The Kolmogorov-Smirnov distance between data and fit is *D* = 0.1, while the log-likelihood ratio between the power law and the exponential fit is R = 295 (*p* < 10^-5^). (B) The distribution of avalanche duration follows a power law with exponent α = 1.973 ±0.002 (fit ±std. error on the fit), followed by an exponential cutoff (pooled data, 10 subjects). The Kolmogorov-Smirnov distance between the data and the fit is *D* = 0.07 The log-likelihood ratio between the power-law and the exponential fit is *R* = 95 (*p* < 10^-5^). Maximum likelihood estimation of the power law exponents were performed using the Powerlaw Python package (Alstott et al., 2014) over the range of values indicated by the thick black lines.

**Figure 3:**
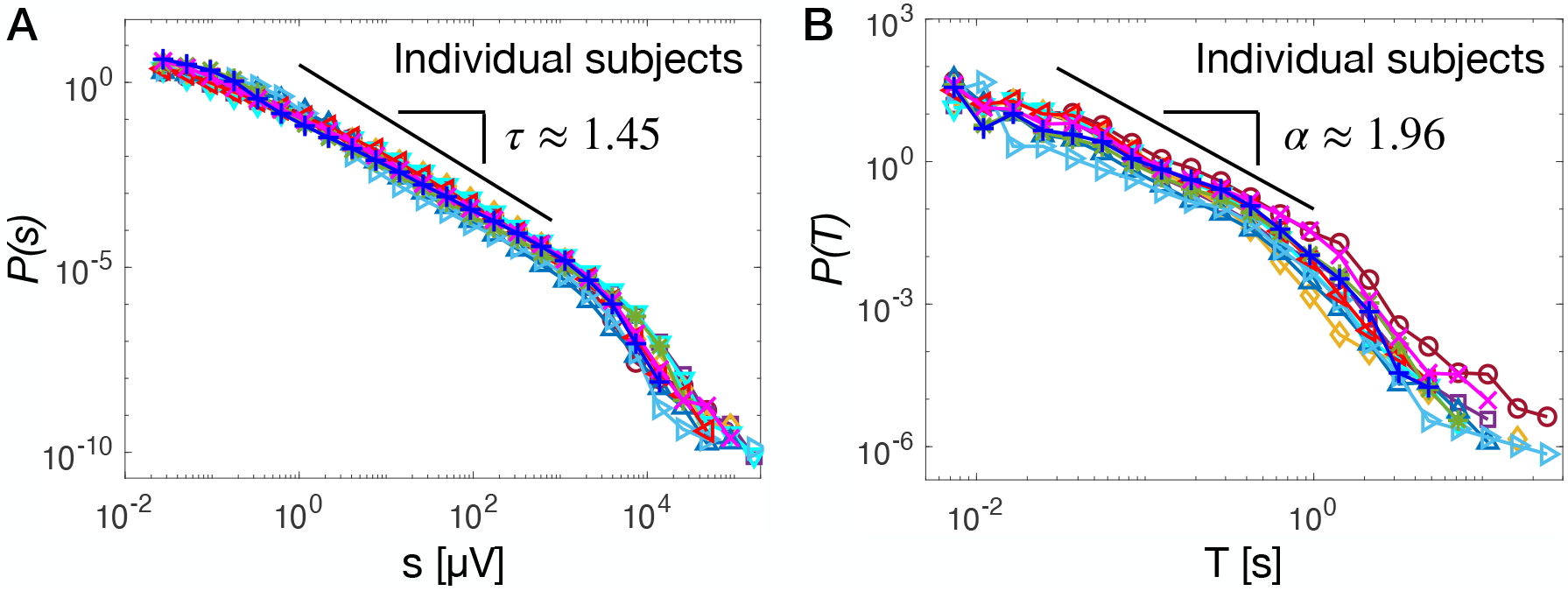
Avalanche size and duration distributions consistently follow a power law behavior during sleep periods across individual subjects. (A) The distribution of avalanche sizes follows a power law with exponent *τ* = 1.45 ±0.09 (mean ±SD). The power law regime is followed by an exponential cut off in all individual subjects. (B) The distribution of avalanche duration follows a power law with exponent *τ* = 1.96±0.16 (mean ±SD), followed by an exponential cutoff. For each individual subject, maximum likelihood estimation of the power law exponents were performed over the range of values corresponding to the thick black line using the Powerlaw Python package (Alstott et al., 2014).

**Figure 4:**
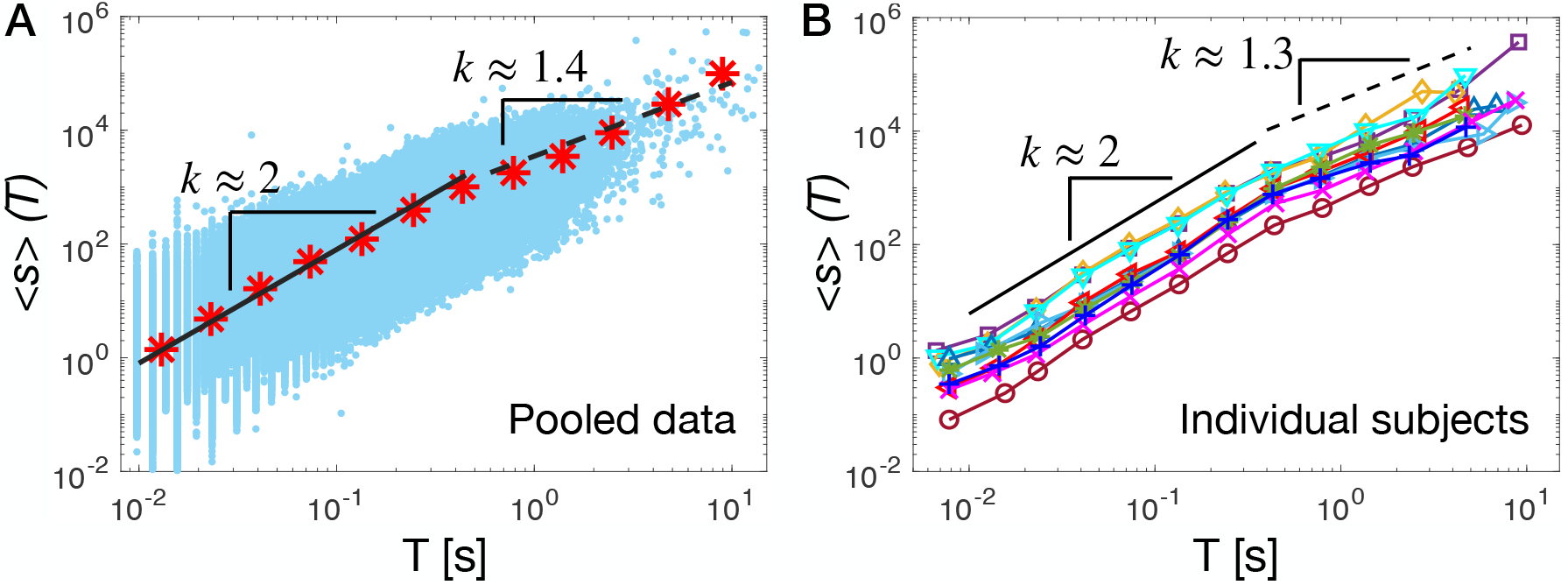
Avalanche sizes and durations are connected by the scaling relationship 〈*s*〉 ∝ *T^k^* consistent with underlying criticality. (A) Average avalanche size as a function of the avalanche duration **T** (red stars; pooled data, 10 subjects). The average avalanche size scales as 〈*s*〉 (*T*) ∝ *T* ^*k*^ with *k* = 1.89 for *T*’s within the scaling regime of the distribution *P*(*T*). This power-law regime is followed by a crossover to a power-law with a significantly smaller exponent *k* =1.4 for larger *T*’s. The thick black line is a power law fit for 0.01 < *T* < 0.4 s; dashed black line is a power law fit for 0.4 < *T* ≤ 5. Blue dots: (s,T) scatter plot. (B) Average avalanche size as a function of the avalanche duration *T* for all individual subjects. The relationship between avalanche sizes and durations is consistent across subjects, showing a crossover from an exponent *k* = 1.96 ± 0.13, and *k*_1_ = 1.32 ± 0.19 (mean ± SD).

An avalanche was defined as a continuous time interval in which there is at least one excursion beyond threshold in at least one EEG channel (Fig. 1). Avalanches are preceded and followed by time intervals with no excursions beyond threshold on any EEG channel (Beggs and Plenz, 2003; Meisel et al., 2013). The size of an avalanche, *s*, was defined as the sum over all channels of the absolute values of the signals exceeding the threshold.

To characterize the relationship between the avalanche dynamics and the sleep macro-architecture, we calculate for each subject the avalanche density as a function of time, i.e. the fraction of time occupied by avalanches, measured as

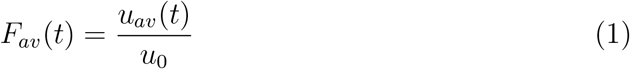

where *u_av_*(*t*) is the amount of time occupied by avalanches in a sliding window of length T (sliding step = 1/(sampling frequency)), and *u_0_* = *T*. The window length T has been chosen equal to 10 seconds (*T* = 10 s) as this is the order of magnitude of the largest avalanches in our recordings.

To characterize the relationship between the avalanche dynamics and the sleep micro-architecture, we compute the Pearson correlation coefficient between the avalanche occurrence and the CAP measures, on a time scale dictated by the sampling rate of the recordings. Given the binary values *x_i_* = 0, 1, *y_i_* = 0, 1, where *x_i_* = 1 indicates the presence in the sample *i* of an ongoing avalanche, and *y_i_* = 1 indicates presence of a particular feature of the CAP framework (CAP, NCAP, subtypes A1, A2, A3, all A phases, phases B), we computes the Pearson correlation coefficient as:

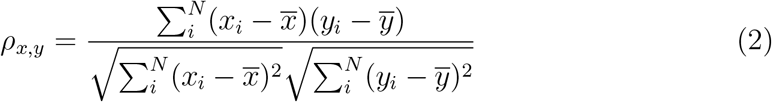

where 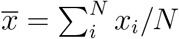 and *N* is the number of samples in the sleep recordings. Being binary values, the Pearson correlation coefficient is equivalent to the *φ* coefficient. The Pearson correlation coefficient has been also evaluated between avalanche occurrence and sleep stages using Eq. 2, where *x_i_* = 1 indicates the presence (*x_i_* =0 absence) in the sample *i* of an ongoing avalanche, and *y_i_* = 1 indicates presence (*y_i_* = 0 absence) of a particular sleep stage (REM,N1,N2,N3).

### 2.4 Statistical analysis

Maximum likelihood estimation of power law exponents for avalanche size and duration distributions was performed using the Power law Python package (Alstott et al., 2014). The power law fit minimized the Kolmogorov-Smirnov distance between original and fitted values, *D* = *sup_x_*|*F_data_*(*x*) – *F_fit_*(*x*)|, where *F_data_* is the empirical cumulative distribution function (CDF) and *F_fit_* the fitted CDF. The power law fit was compared to an exponential fit by evaluating the log-likelihood ratio *R* = *lnL_p_*/*L_e_*, where 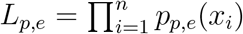 is the likelihood. *R* is positive if the data are more likely to follow a power law distribution, and negative if the data are more likely to follow exponential distribution. The statistical significance for *R* (*p*-value) was estimated in the Power law Python package (Alstott et al., 2014). For further details see (Clauset et al., 2009). Pairwise comparisons in Fig. 5 and 6 were conducted using Students two-tailed t-test performed in Matlab (Mathworks).

**Figure 5:**
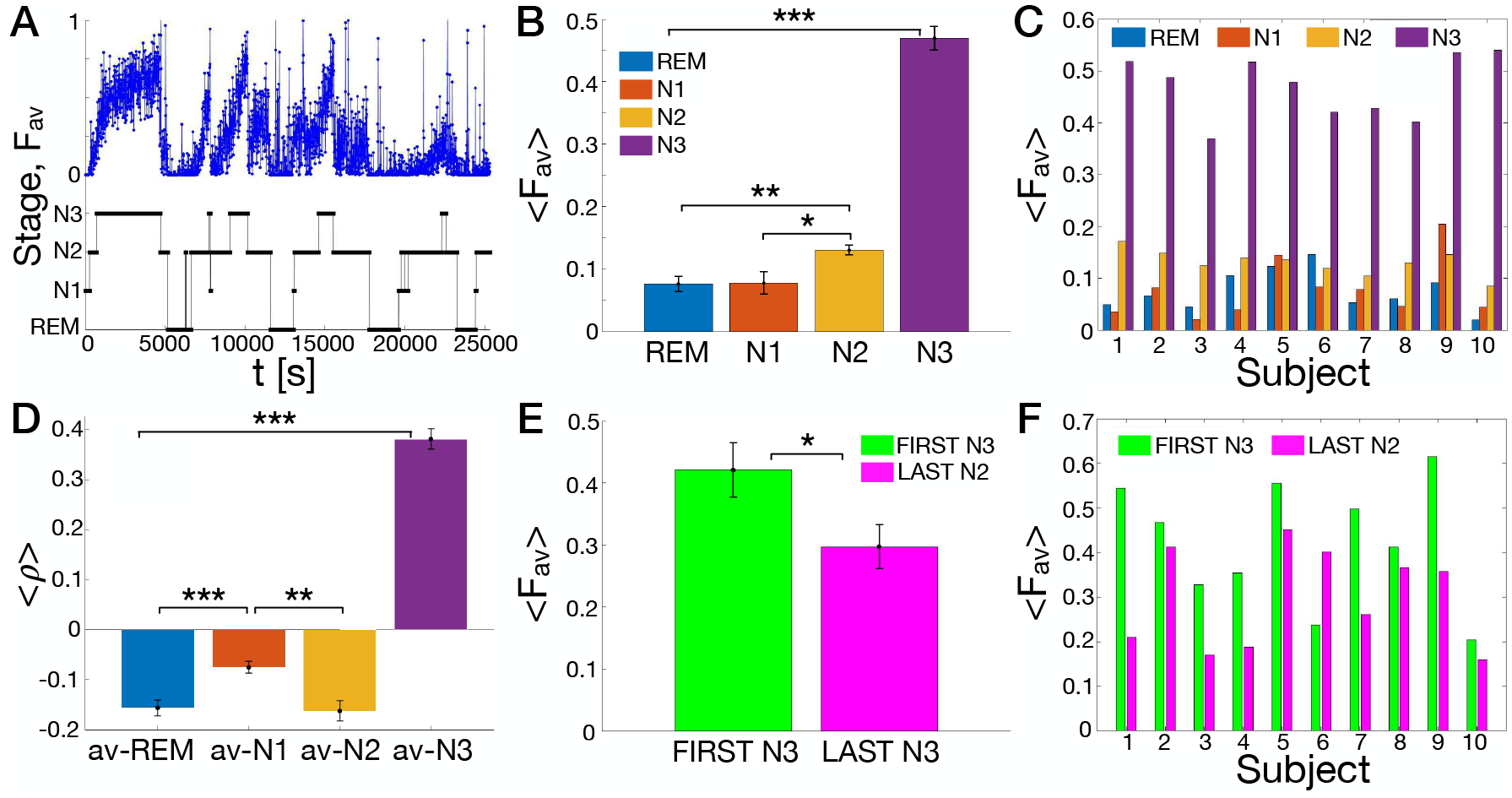
Overnight sleep macro-architecture is associated with strong modulation of avalanche dynamics. (A) The density of avalanches (blue dots), *F_av_*(*t*), is shown as a function of time, together with the corresponding sleep stages and sleep stage transitions (REM, N1, N2, N3 black line) for an individual subject. *F_av_*(*t*) increases gradually in N2 and N3, and then abruptly decreases when transitioning from N3 to either N2, N1 or REM. Waking periods during sleep have been removed. (B) Mean avalanche density for each sleep stage (REM, N1, N2, N3) averaged across subjects. The density *F_av_*(*t*) is highest in N3 and gradually decreases for N2, N1, and REM. Differences between N3 and all other sleep stages are significant (N3 versus N2: *p* = 1.7 · 10^-9^; N3 versus N1: *p* = 1.5 · 10^-11^; N3 versus REM: *p* = 1.6 · 10^-11^). *F_av_* in N2 is significantly different from the density in N1 (*p* = 0.019) and REM (*p* = 0.002). (C) Mean avalanche density for each sleep stage and for each individual subject. The behavior observed for the group average is consistent across individual subject, N3 being the sleep stage with the highest density of avalanches. (D) The mean Pearson correlation coefficients *ρ_x,y_* (see Eq. (2)) between avalanche occurrence and sleep macro-architecture (namely REM,N1,N2,N3) shows that avalanches tend to occur mostly during N3. In all bar plots error bars indicate the standatd error of mean. Differences between N3 and all other sleep stages are significant (N3 versus N2: *p* = 2.7 · 10^-13^; N3 versus N1: *p* = 8.7 · 10^-12^; N3 versus REM: *p* = 1.7 · 10^-13^). N2 is significantly different from N1 (*p* = 0.002), and N1 is significantly different from REM (*p* = 0.0006). (E) Mean density of avalanches in the first and last N3 stage of the recordings averaged over all subjects. We observe that the density is significantly higher during the first N3 (*p* = 0.04). (F) Avalanche density in the first N3 (blue) and last N3 (red) for each individual subject. The density is higher in the first N3 for all subjects but the subject #6, for which we observe that the density is higher in the last N3. Such deviation from the average behavior may be related to general differences we observed in sleep of subject #6. For instance, this subject presented an unusually short duration of the N3 stage at the beginning of the night, followed by a gradual increase of N3 in the second half of the sleep. Significance legend: *** for *p* < 0.001; ** for *p* < 0.01; * for *p* < 0.05. The *** in panel B and D refers to the pairwise comparison between N3 and all the other sleep stages. The ** in panel B refers to the pairwise comparison between N2, N1, and REM. Differences are not significant where no stars are reported.

**Figure 6:**
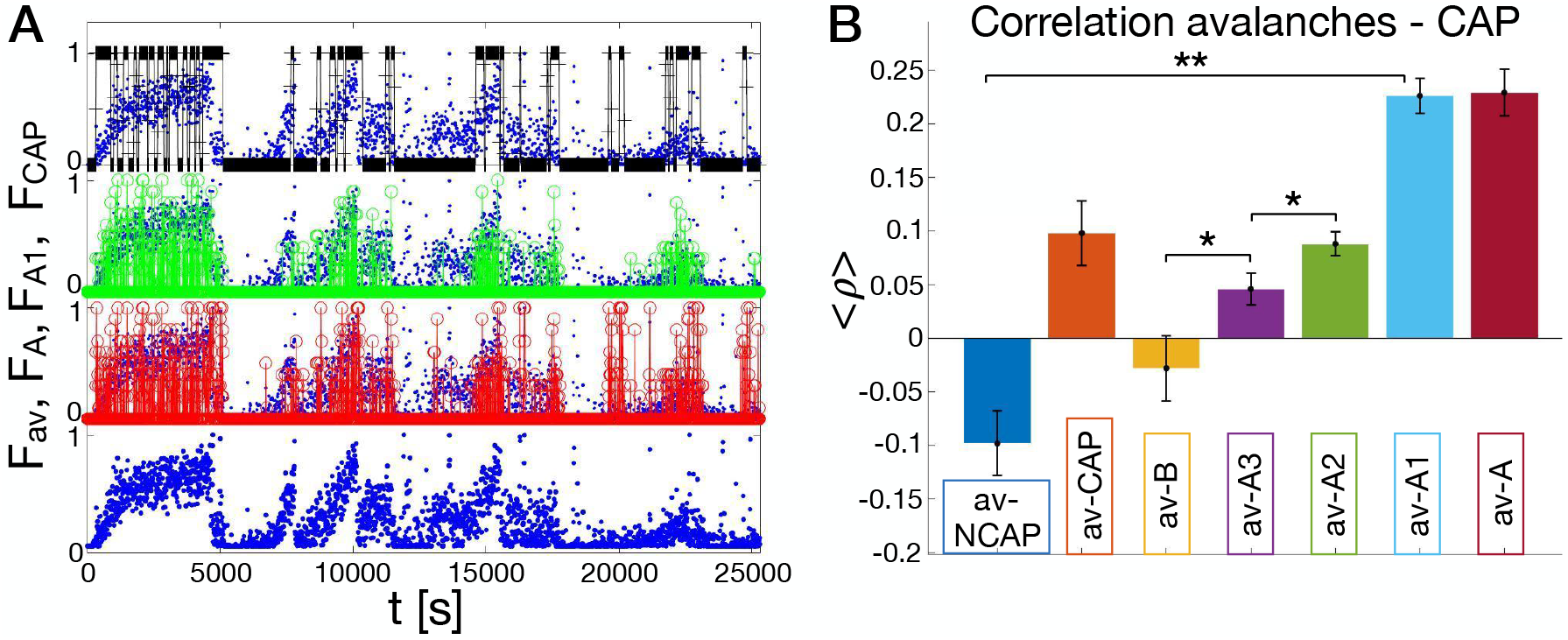
Occurrence of neuronal avalanches is coupled with the occurrence of the CAP. (A) Density of avalanches versus density of CAP phases as function of time for an individual subject. Density of avalanches in blue, density of phase A in red, density of phase A1 in green, density of CAP in black. (B) The mean Pearson correlation coefficients *ρ*(*x, y*) (average over subjects; see Materials and Methods, Eq. (2)) between avalanche occurrence and micro-architecture features (NCAP, CAP, B, A, and A subtypes A3,A2,A1). Error bars indicate the standard error of the mean. Differences are all significant (*p* < 0.01 for all couples except av-B versus av-A3 and av-A2 versus av-A3) but av-CAP versus av-A3 and av-A2, av-A versus av-A1, and av-NCAP versus av-B. Significance legend: *** for *p* < 0.001; ** for *p* < 0.01; * for *p* <0.05. The ** in panel B refers to the pairwise comparison between av-A1 and all the other bars but av-A. Differences are not significant where no stars are reported.

## 3 Results

### 3.1 Critical exponents and scaling relations for neuronal avalanches during sleep

To characterize cortical dynamics underlying sleep macro-architecture and sleep micro-architecture, we identify neuronal avalanches and investigate signatures of criticality across the entire sleep period. To this end, we compute the distribution of avalanche sizes, *P*(*s*), and avalanche durations, *P*(*T*). In Fig. 2 we show the distributions *P*(*s*) and *P*(*T*) for all subjects. We find that both the size and duration distributions are well described by a power law, *P*(*s*) ∝ *s^−τ^* and *P*(*T*) ∝*T^−α^*, respectively. In both distributions the power law regime is followed by an exponential cutoff (Fig. 2). Power laws are the hallmark of criticality, and imply absence of characteristic scales in the underlying dynamics (Stanley, 1971). In this context, the observed power law distributions indicate that neuronal avalanches have no characteristic size and duration, namely they are scale-free. Our analysis shows that the exponent *τ* for the size distribution is close to 3/2 (*τ* = 1.438 ±0.001) (fit ±error on the fit), while the exponent *α* for the duration distribution is close to 2 (1.973 ±0.002). We compared the power law with an exponential fit by evaluating the log-likelihood ratio 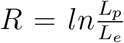 between the likelihood *L_p_* for the power law and *L_e_* for the exponential fit (Materials and Methods). We found *R* = 295 for the size and *R* = 95 (*p*-value < 10^-5^; see Materials and Methods) for the duration distribution, indicating that the respective power laws better describe the empirical distributions. Importantly, we observe that the power law exponents *τ* and *α* are robust and weakly depend on the scale of analysis (Fig. S1)—e.g. the threshold used to identify avalanches——, and are consistent across subjects (Fig. 3). In Fig. 3 we show the avalanche size and duration distributions for all individual subjects. Both distributions show little variability across subjects, and follow a power law with exponents *τ* = 1.45 ±0.09 and *α* = 1.96 ± 0.16 (mean ±SD), in agreement with values measured on pooled distributions (Fig. 2). We note that this values are fully consistent with the the values predicted within the mean-field directed percolation (MF-DP) universality class—3/2 and 2, respectively (Pruessner, 2012).

Next, we analyze the relationship between avalanche sizes and durations. Near criticality the average avalanche size 〈*s*〉 is expected to scale as a power of the duration *T*, namely 〈*s*〉 ∝ *T^k^* (Pruessner, 2012). We find that such a power law relationship between avalanche sizes and durations holds during sleep (Fig. 4). In particular, we observe that, for *T*’s smaller than the duration corresponding to the onset of the exponential cutoff in the distribution *P*(*T*) (Fig. 2 and 3), the average size scales as 〈*s*〉) ∝ *T^k^* with *k* ≈ 2 (Fig. 4). For larger durations, we observe a crossover to a power law relationship with a smaller exponent *k* ≈ 1.3 (Fig. 4). Importantly, the exponent k is robust and independent of the threshold *θ* used to detect neuronal avalanches (Fig. S1). Moreover, we observe that the relation 〈*s*〉 ∝ *T^k^* is consistent across individual subjects (Fig. 4B), the exponent *k* showing little variability across subjects. Specifically, we find *k* = 1.96 ± 0.13 (mean ± SD) for *T*’s smaller than the duration corresponding to the onset of the exponential cutoff in the distribution *P*(*T*), and *k* = 1.32 ± 0.19 (mean ± SD) for larger T’s (Fig. 4B).

Notably, we find that the exponent *k* measured in Fig. 4 is in agreement, within errors, with the value predicted by the scaling relation

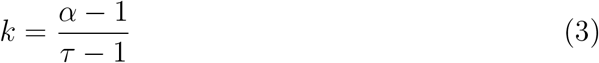

in the context of crackling noise (Sethna et al., 2001). Indeed, we have that 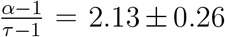 (mean ±std error), and *k* = 1.96 ± 0.04 (mean ± std error). The scaling relation in Eq. 3 has a general validity in avalanche dynamics, as shown in (Scarpetta et al., 2018; Fosque et al., 2021), where Eq. 3 was derived with the only hypothesis that *P*(*s*) ∝ *s*^-τ^ and *P*(*T*) ∝ *T^−α^*, and that the size fluctuations for fixed durations are small and can be neglected.

In sum, during sleep, the values of the critical exponents *τ*, *α* and *k* are very close to the ones predicted for the critical branching process, i.e. the mean field directed percolation (MF-DP) universality class, with exponents *τ* = 3/2 for the size and *α* = 2 for the duration distribution, and *k* = 2 (Pruessner, 2012).

### 3.2 Avalanche dynamics and sleep macro-architecture

We have shown that, during sleep, neuronal avalanches are characterized by a robust scaling behavior in their size and duration distributions (Fig. 2 and 3), and that avalanche size and duration are linked by precise scaling relationships (Eq. 3 and Fig. 4). These observations are robust and consistent across subjects, and indicate underlying tuning to criticality during sleep. Next, we investigate the relationship between critical avalanche dynamics, sleep stages, and sleep stage transitions.

We first characterize sleep macro-architecture across all subjects. The main sleep parameters are described in Table 1 (macro-structural measures). The average TST across the 10 subjects was 423.9 min, with a mean SE of 88.92%. Around 56% of TST was spent in light sleep (N1 = 7.23%, N2 = 48.47%), 23.99% in deep sleep (N3 = 23.99%), and 20.30% in REM.

**Table 1:** Average characteristics of sleep macro-architecture across the analyzed subjects (n = 10). For each measure mean and standard deviation (SD) are reported. SE = sleep efficiency (SE), TST= total sleep time, WASO = Wake After Sleep Onset.

| Measure | MEAN | SD |
| --- | --- | --- |
| Sleep latency (min) | 9,90 | 12,25 |
| SE (%) | 88,92 | 9,28 |
| TST (min) | 423,90 | 63,31 |
| WASO (min) | 40,97 | 30,46 |
| Stage N1 (min) | 28,75 | 17,47 |
| Stage N1 (%) | 7,23 | 5,20 |
| Stage N2 (min) | 207,10 | 50,78 |
| Stage N2(%) | 48,47 | 6,67 |
| Stage N3 (min) | 99,35 | 13,40 |
| Stage N3(%) | 23,99 | 4,90 |
| NREM sleep (min) | 335,20 | 41,67 |
| REM sleep (min) | 88,65 | 32,02 |
| REM sleep (%) | 20,30 | 5,14 |

To study the interplay between sleep macro-architecture and avalanche dynamics, we introduce the avalanche density, *F_av_* (t), defined as the amount of time occupied by avalanches in a sliding window of length *u*_0_ (Materials and Methods), and study the temporal evolution of *F_av_*(*t*) in relation to the sleep macro-architecture. In the following we fix *u*_0_ = 10 s, which approximately corresponds to the largest avalanche duration we observed (Fig. 2 and 3). In Fig. 5A we show the avalanche density *F_av_*(t) as a function of time for an individual subject, together with the corresponding hypnogram. We observe that *F_av_*(*t*) gradually increases in parallel with sleep deepening, i.e. going from REM to N1, N2, and finally N3: *F_av_* is very small during stage N1, reaches an intermediate value during stage N2, and increases substantially during stage N3, where it peaks slightly before the following transition back to N2 and REM (Fig. 5A). Although the avalanche density tends to decrease across the night and is, on average, much smaller at the end of the night, we find that this trend repeats throughout the night in correspondence to the descending REM → N3 of the NREM-REM sleep cycle. In contrast to this gradually increasing trend, we observe that the avalanche density decreases rather abruptly with transitions from N3 to N2 and N1—the ascending phase of the NREM-REM sleep cycle. In sum, we find that the avalanche density gradually increases during the descending slope of each sleep cycle, whilst it rapidly decreases in the ascending slope of the same cycles that precedes the onset of REM sleep (Fig. 5A).

Our analysis shows that the density of avalanches is significantly higher during N3 as compared to N2, N1, and REM (Fig. 5B, C). The analysis of the Pearson correlation coefficient *ρ_x,y_* (Materials and Methods, Eq. 2) shows that avalanche occurrence, on average, is positively correlated with N3, while it is either weakly or slightly negatively correlated with other sleep stages (Fig. 5D). Finally, we observe that, during N3, the avalanche density tends to increase with time (Fig. 5A). This suggests that the mechanisms related to generation of neuronal avalanches become more and more effective during SWS, and move the system towards the deepest phase of sleep.

Importantly, we notice that the avalanche density peak—typically located within N3 periods—is higher in the first half of the night, progressively decreases during the second half of the night. To quantify the significance of this behavior with respect to the characteristics of neuronal avalanches, we compare the avalanche density, as well as avalanche size and duration distributions, in the first and last N3 stage of the sleep recordings. We find that avalanche size and duration distributions in the first N3 are comparable to the distributions calculated in the last N3 (SI, Fig. S2). Furthermore, the scaling relation 〈*s*〉 ∝ *T^k^* between avalanche size and duration is satisfied both in the first and last N3, with the same values of the exponent *k* (SI, Fig. S1). On the other hand, we observe that the avalanche density is significantly higher during the first N3 as compared to the last N3 (Fig. 5E, F) (*t*-test: *p* = 0.04).

This is consistent across subjects (Fig. 5F), with only one exception (subject #6, Fig. 5F).

### 3.3 Avalanche dynamics and sleep micro-architecture

The analysis of the avalanche density across sleep stages has shown that neuronal avalanches tend to occur with higher frequency during NREM sleep. However, NREM sleep has a complex micro-architecture that is characterized by the CAP phenomenon (Terzano et al., 2002). In our data, the mean CAP rate was 49.19% with the following distribution across NREM stages: N1= 41.69%, N2= 48.36%, and N3= 53.37% (Table 2). On average, subjects presented 37.1 CAP sequences per night, with a mean duration of 4.55 min. With respect to CAP subtypes distribution, 206 were A1 (25.7% of the CAP time); 67.2 were A2 (9.2% of the CAP time), and 83.8 were A3 (14.19% of the CAP time). A1’s were more present during stage N3 (50.21%) as compared to N2 (5.72%) and N1 (1.49%), in agreement with previous studies (Halász et al., 2004). On the other hand, subtypes A2 and A3 predominated in stage N1 (particularly A3, 37,77%) and N2 (14.39% for A2 and 17.46% for A3) 2.

**Table 2:** Average characteristics of sleep micro-architecture across the analyzed subjects (n = 10).

| **Measure** | **MEAN** | **SD** |
| --- | --- | --- |
| **CAP time (minutes)** | 162,51 | 42,3 |
| **CAP rate (%)** | 49,05 | 14,4 |
| **CAP sequences (n)** | 37,1 | 8,3 |
| **CAP sequences length (min)** | 4,55 | 1,6 |
| **CAP cycle (n)** | 357,6 | 104,1 |
| **Phase A length (s)** | 8,59 | 1,4 |
| **Phase B length (s)** | 20,67 | 3,5 |
| **Phase A1 (n)** | 206 | 86,9 |
| **Phase A2 (n)** | 67,2 | 39,2 |
| **Phase A3 (n)** | 83,8 | 34,9 |
| **CAP A1 Rate (%)** | 25,7 | 11,7 |
| **CAP A2 Rate (%)** | 9,2 | 5,8 |
| **CAP A3 Rate (%)** | 14,19 | 7,4 |
| **CAP Rate N1 (%)** | 41,69 | 17,3 |
| **CAP Rate N2 (%)** | 48,36 | 19,3 |
| **CAP Rate N3 (%)** | 53,37 | 18,8 |
| **CAP A1 N1 (%)** | 1,75 | 3,0 |
| **CAP A1 N2 (%)** | 16,9 | 12,2 |
| **CAP A1 N3 (%)** | 50,21 | 19,5 |
| **CAP A2 N1 (%)** | 2,37 | 3,1 |
| **CAP A2 N2 (%)** | 14,39 | 9,4 |
| **CAP A2 N3 (%)** | 5,72 | 2,1 |
| **CAP A3 N1 (%)** | 37,77 | 17,9 |
| **CAP A3 N2 (%)** | 17,46 | 9,4 |
| **CAP A3 N3 (%)** | 1,49 | 1,7 |
| **Subtype A1 duration (s)** | 6,42 | 2,0 |
| **Subtype A2 duration (s)** | 8,63 | 2,0 |
| **Subtype A3 duration (s)** | 12,72 | 1,32 |

To dissect the relationship between CAP and occurrence of neuronal avalanches during NREM sleep, we compare the time course of the avalanche density with the density of distinct CAP phases (Fig. 6A) defined as *F_X_* (*t*) = (*u_X_* (*t*))/*u*_0_, where X denotes the specific CAP phase—A, A1, A2, A3, B—and *u_X_*(*t*) the time occupied by the specific CAP phase in a window of length *u*_0_= 10 s. We observe a remarkable time correspondence between the temporal profile of the density of avalanches *F_av_*(*t*) and the density of CAP, with the peaks in avalanche density corresponding to high density of CAP—in particular phase A and A1 (Fig. 6A). Specifically, we notice that, with sleep deepening, the progressive increase of CAP density is accompanied by a parallel increase in avalanche density. We find that the percentage of phase A occupied by neuronal avalanches is about 42.16%, while the percentage of sleep time occupied by avalanches is 19,21% (Materials and Methods). Interestingly, CAP phase A1 is even richer in avalanches compared to CAP A phases A2 and A3 (53,32% versus 43,84% and 27,72%, respectively).

The physiological increase of CAP cycles during N2 and N3, indirectly leads to a reduction of time occupied by NCAP sleep. Furthermore, during the deepest stages of NREM sleep, CAP’s typically present shorter phases B. These changes in the sleep micro-dynamics lastly sustain the observed increase of avalanche density.

Next, we measure the Pearson correlation coefficients between occurrence of neuronal avalanches and different CAP phases (see Materials and Methods, Eq. 2). We find positive correlations between occurrence of avalanches and CAP phase A, in particular CAP phase A1 (Fig. 6B). On the contrary, we observe negative correlations between occurrence of avalanches, CAP phase B, NCAP periods. This indicates that the occurrence of avalanches during NREM sleep is strictly related to occurrence of CAP, and in particular CAP phase A1. These results are consistent across subjects, as shown in Table 3.

**Table 3:** Pearson correlation coefficient between avalanche occurrence and CAP subtypes for the analyzed individual subjects (n = 10) (Materials and Methods).

| Subject | av-NCAP | av-CAP | av-B | av-A3 | av-A2 | av-A1 | av-A |
| --- | --- | --- | --- | --- | --- | --- | --- |
| #1 | -0,240 | 0,240 | 0,113 | 0,050 | 0,080 | 0,270 | 0,260 |
| #2 | -0,140 | 0,140 | -0,016 | 0,130 | 0,150 | 0,220 | 0,290 |
| #3 | -0,060 | 0,060 | -0,041 | 0,090 | 0,090 | 0,170 | 0,190 |
| #4 | -0,140 | 0,140 | 0,039 | 0,010 | 0,110 | 0,230 | 0,190 |
| #5 | -0,040 | 0,040 | -0,013 | 0,000 | 0,050 | 0,190 | 0,130 |
| #6 | -0,020 | 0,020 | -0,206 | 0,040 | 0,090 | 0,250 | 0,270 |
| #7 | -0,070 | 0,070 | -0,074 | 0,080 | 0,110 | 0,200 | 0,240 |
| #8 | -0,220 | 0,220 | 0,056 | 0,040 | 0,110 | 0,330 | 0,340 |
| #9 | -0,140 | 0,140 | 0,016 | 0,050 | 0,060 | 0,240 | 0,230 |
| #10 | 0,080 | -0,080 | -0,158 | -0,030 | 0,030 | 0,160 | 0,130 |
| **Mean** | -0,099 | 0,099 | -0,029 | 0,046 | 0,088 | 0,226 | 0,227 |
| **Std error** | 0,030 | 0,030 | 0,097 | 0,219 | 0,016 | 0,011 | 0,015 |

## 4 Discussion

In this paper we analyzed the scaling properties of neuronal avalanches during sleep in healthy volunteers, and investigated the relationship between avalanche dynamics and sleep macro- and micro-architecture, with a particular focus on the cyclic alternating patterns (CAP). We showed that the scaling exponents characterizing neuronal avalanches are consistent with the MF-DP universality class, and obey the scaling relations theoretically predicted. This indicates that, during physiological sleep, brain dynamics is consistent with criticality and is satisfactorily described by the MF-DP universality class. Furthermore, we introduced a measure—the density of avalanches—to quantify the relationship between avalanche dynamics and sleep macro- and micro-architecture. Our analysis showed that distributions of avalanches in time is not random but closely follow the descending and ascending phase of the NREM-REM cycles. Within such cycles, the presence of neuronal avalanches is linked to the occurrence of CAP during NREM sleep. Specifically, we found that the density of avalanches is higher during NREM, and, within NREM sleep, avalanche occurrence is positively correlated with the phase A of the CAP, in particular the phase A1. This suggests a close relationship between modulation and control of brain criticality, sleep macro- and micro-architecture, and brain function, which we discuss in turn.

### Brain dynamics and criticality during sleep

Empirical evidence indicates that the human brain operates close to a critical regime both in resting wakefulness and during sleep (Priesemann et al., 2013; Bocaccio et al., 2019; Allegrini et al., 2015; Lombardi et al., 2021b, 2020a; Wang et al., 2019). In particular, recent studies suggest that criticality plays a key role in determining the temporal organization of sleep stage and arousal transitions (Lombardi et al., 2020a; Wang et al., 2019). However, critical dynamics during sleep remains poorly understood. In this respect, a key open question concerns the universality class to which brain criticality obeyed during sleep. To the best of our knowledge, this is the first study investigating this problem, and exploring the scaling relation among critical exponents of neuronal avalanches during sleep. We reported a picture that is fully consistent with the MF-DP universality class. Indeed, we have shown that (i) the critical exponents for the avalanche size and duration distributions are very close to the prediction of the critical branching process, MF-DP universality class, i.e. *τ* = 3/2, *α* = 2, respectively; (ii) the exponent *k* connecting sizes and durations is very close to 2, as predicted; (iii) the exponents *τ*, α, and *k* correctly satisfy the expected scaling relation.

The exponent *k* has been previously measured in the awake resting-state, from Zebrafish and rats to monkeys and humans (Ponce-Alvarez et al., 2018; Miller et al., 2019; Fontenele et al., 2019; Lombardi et al., 2021a; Mariani et al., 2021; Dalla Porta and Copelli, 2019). In line with our findings, Miller et al. (Miller et al., 2019) found that, in awake monkeys, *k* ≃ 2 in the range corresponding to the power law regime of the size and duration distributions, while *k* ≃ 1 – 1.5 in the region that corresponds to the exponential cut-off of the distributions—where we found *k* ≈ 1.3. Similar results were found in Zebrafish (Ponce-Alvarez et al., 2018). Deviation from the value *k* = 2 was observed in the resting-state of the human brain (Lombardi et al., 2021a), in ex-vivo turtle visual cortex (Shew et al., 2015), in the barrel cortex of anesthetized rats (Mariani et al., 2021), in cortex slice cultures (Friedman et al., 2012), and in freely behaving and anesthetized rats (Fontenele et al., 2019). Notably, a recent work (Apicella et al., 2022) has shown that, in a 2D neural network, the value of the exponent *k* is related to the network connectivity, with *k* ≃ 1.3 for a 2D connectivity, and *k* = 2 when the mean-field approximation is justified, namely when the spatial extension can be considered small as compared to the system’s connectivity range. This suggests that the crossover observed in *S*(*T*) (Fig. 4) from *k* ≃ 2 to *k* ≃ = 1.3 could be due to the different nature of small, localized avalanches, which propagate over a densely connected network, and larger avalanches, which rely on the structured topology of large scale brain networks with sparser and long-range connections. Subsampling in brain activity recordings has also been suggested as a potential origin of the observed scaling exponents (Carvalho et al., 2021).

### Neuronal avalanches and sleep macro-architecture

The static properties of neuronal avalanches during sleep, i.e. size and duration distributions, have been investigated in previous studies. Analyses of scalp EEG and human intracranial depth recordings showed that such distributions follow a similar power law behavior across the sleep-wake cycle, with exponents in line with our observations (Allegrini et al., 2015; Priesemann et al., 2013). Similarly, the analysis of whole-brain fMRI data confirmed a robust critical (or near-critical) behavior from wakefulness to deep sleep, with little differences in the power-law exponent of the avalanche size distribution (in particular between wakefulness and stage N2) (Bocaccio et al., 2019).

On the other hand, here we have shown that, although the static properties remain fairly stable across different sleep stages (Bocaccio et al., 2019; Allegrini et al., 2015; Priesemann et al., 2013), avalanche dynamics is modulated by the ascending and descending slope of the NREM-REM sleep cycles. By analyzing the temporal evolution of the avalanche density, we found that avalanche occurrence markedly and progressively increases with NREM sleep stages N2 and N3 and, specifically, during periods of sleep deepening (descending slope of sleep cycles), in parallel with the increase of SWA. On the contrary, the abrupt decrease in avalanche density during the ascending slope of sleep cycles suggests a negative influence from REM-on/wakefulness circuits with respect to their appearance. The different behavior of avalanche density during the descending and ascending slopes of the sleep cycles was not previously observed, despite the crucial role of such dynamics for sleep regulation. In terms of sleep physiology, the descending and ascending slopes of sleep cycles are markedly different: during the descending slope, sleep-promoting forces are stronger, the thalamo-cortical system works in the burst-firing mode and brainstem cholinergic pathways are tonically repressed. Conversely, during the ascending slope, the NREM driving forces become weaker, sleep is more vulnerable towards pro-arousal intrusions and REM-promoting outputs prevail (Halász et al., 2004). Taking this into account, our results suggest that avalanche occurrence is not random across the sleep cycles, but instead contributes to define and sustain the dynamical interplay between sleep-wake promoting networks.

### Avalanches and sleep micro-architecture

Sleep architecture is composed of numerous oscillatory patterns, including, above all, the CAP (Terzano et al., 2000). CAP’s occur on time scales of seconds or minutes, accompany sleep stage shifts, and contribute to the organization of sleep cycles. The CAP is a periodic EEG activity that reflects a state of brain instability, and is characterized by the alternation of phases of higher EEG amplitude (CAP phases A, “activation phases”) separated by periods of lower EEG amplitude (CAP phases B, “de-activation phases”)—both phases lasting between 2 and 60 seconds. Conversely, the NCAP is defined as a period of sustained physiologic stability. CAP phases A can be further subdivided into three subtypes: A1, A2 and A3. Isolated A1 phases, not followed by a subsequent phase A within 60 seconds, are scored as NCAP, confirming that the dynamic interplay between phases of activation/baseline is key characteristic of the CAP framework.

Our analyses demonstrated positive correlations between CAP and avalanche occurrence, and negative correlations for NCAP sleep. Such link suggests a close relationship between CAP and brain tuning to criticality during sleep, a key aspect that should be further investigated in future work.

Although the definition of avalanches (large, collective non-gaussian fluctuations of brain activity) is not related to the definition of CAP phase A, our results show that neuronal avalanches are correlated with the occurrence of CAP phase A. In particular, we observed stronger correlations between avalanche occurrence and the CAP A1 subtype, and weaker positive correlation with subtypes A2 and A3. Interestingly, the correlation between avalanches and the phase A of the CAP is more prominent than the correlation with the CAP itself— phase A and phase B together. We speculate that this could be due to the opposite significance of CAP phase A and B with respect to sleep dynamics. Electrophysiologically the phase B is characterized by the rebound of background EEG activity after the strong ‘activation’ driven by the phase A. Compared to phase A, the phase B could be described as “lower arousal reaction” or vehicles of deactivation (Parrino et al., 2012). Importantly, we did not observe significant correlation between avalanche occurrence and phase B, corroborating our assumption about the relationship between CAP phase A and avalanches. The prominent correlation between avalanche occurrence and CAP “activation phase” A1 may suggest that neuronal avalanches emerge at the edge of a synchronization phase transition, as recent numerical studies indicate (Di Santo et al., 2018; Scarpetta and de Candia, 2014; Scarpetta et al., 2013).

Finally, we note that CAP-A1 physiologically prevail in the first half of the night and during the descending slope of each sleep cycle, boosting or maintaining SWS. Similarly, the avalanche density decreases moving from the first to the last sleep cycle. Hence, both CAP phase A and neuronal avalanches follow a physiological, homeostatic decay throughout the night, and they may both contribute to the build-up of the deepest stages of NREM sleep.

### Neuronal avalanches, CAP, and learning mechanisms: an intriguing hypothesis

Sleep is crucial to renormalize synaptic weight, ensure an optimal and effective network state for information processing, and preserve cognition (Cirelli and Tononi, 2021). Renormalization of synaptic weights taking place during sleep may serve to keep the network close to criticality (Pearlmutter and Houghton, 2009). In line with this view, the here reported higher concentration of avalanches during SWS and CAP-A1 indicate that these states may exert a pivotal role in modulating and restoring brain criticality. Furthermore, because CAP-A1 has been proposed to play a role in the sleep-dependent learning processes (Ferri et al., 2008), our observations point to a functional link between critical avalanche dynamics and sleep-dependent learning processes, as shown in recent numerical studies (Scarpetta and de Candia, 2014; Scarpetta, 2019). Specifically, it has been demonstrated that, within the alternation of up- and down-states observed during SWS, the sequence of avalanches occurring in the up-states correspond to an intermittent reactivation of stored spatiotemporal patterns, a mechanism that is key for memory consolidation (Dupret et al., 2010).

### Conclusions and limitations of the study

Overall, our findings open a novel perspective on the relationship between critical brain dynamics and physiological sleep. We provided a comprehensive account of the critical exponents and scaling relations for neuronal avalanches, demonstrating that brain dynamics during sleep follows the MF-DP universality class. This sets the bases for future investigation of neural collective behaviors occurring during sleep, including their functional role in relation to criticality. As a first step in this direction, our study provides evidence of a functional link between avalanche occurrence, slow-wave sleep dynamics, sleep stage transitions and occurrence of CAP phase A during NREM sleep. As CAP is considered one of the major guardians of NREM sleep that allows the brain to react dynamically to any external perturbation and contributes to the cognitive consolidation processes occurring in sleep, our observations suggest that neuronal avalanches at criticality might be associated with flexible response to external inputs and to cognitive processes—a key assumption of the critical brain hypothesis. This is a crucial aspect that should be investigated in future work. Moreover, based on our results, one could speculate that a relationship between occurrence of neuronal avalanches and physiological sleep measures exists. To address this point, additional studies in pathological sleep conditions where both CAP and criticality-based metrics show a deviation from the physiological parameters are needed (Parrino and Vaudano, 2017; Zimmern, 2020). Future work should also overcome some limitations we acknowledge in the current study. The limited number of subjects and the use of scalp EEG (we enrolled healthy volunteers for which more invasive techniques are not allowed), which limits the analysis of collective neural dynamics

## Acknowledgments

FL acknowledges support from the European Union’s Horizon 2020 research and innovation program under the Marie Sklodowska-Curie Grant Agreement No. 754411, and from the Austrian Science Fund (FWF) under the Lise Meitner fellowship No. PT1013M03318. IA acknowledges financial support from the MIUR PRIN 2017WZFTZP.

## Supplementary Information

**Fig. S1:**
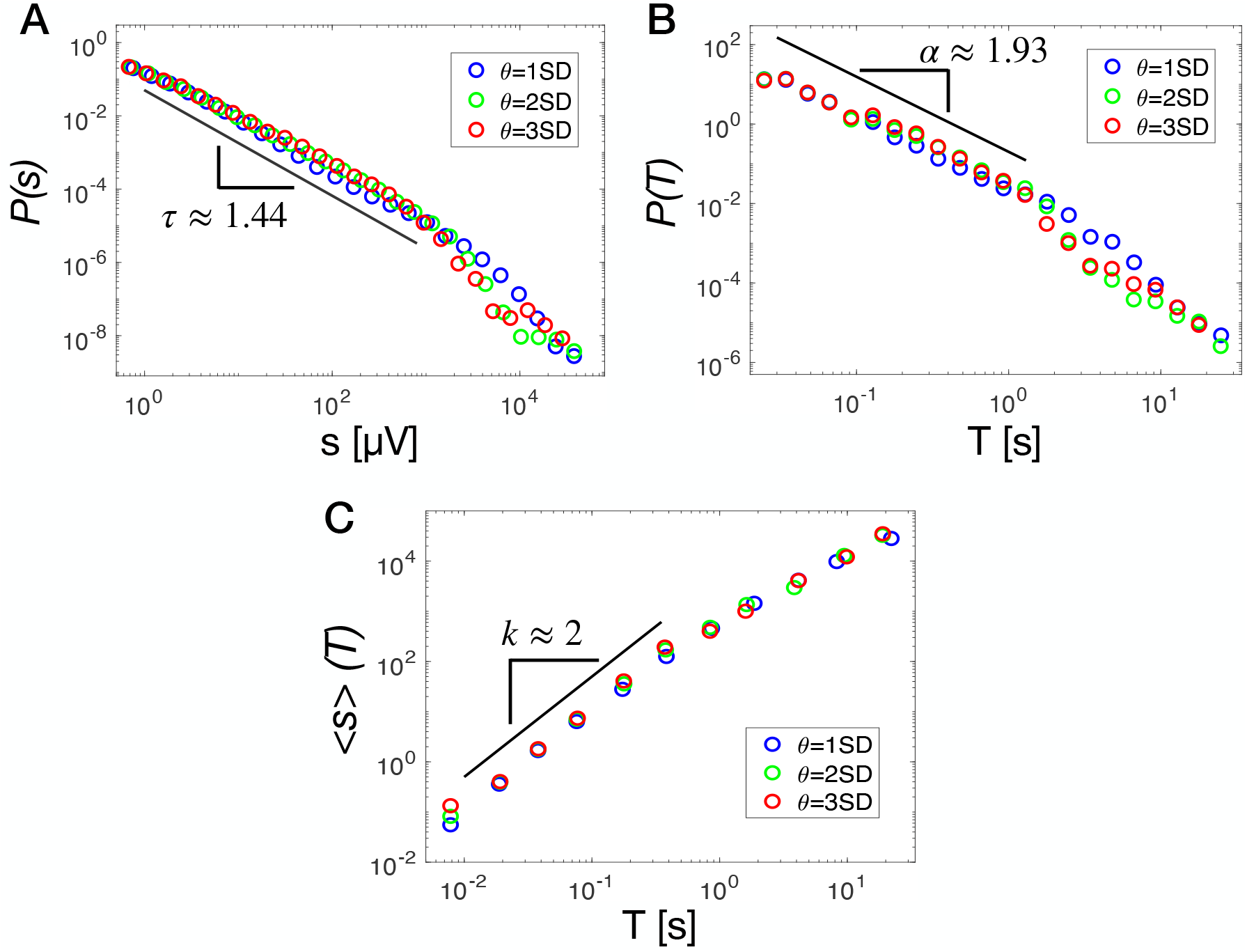
Avalanche size (A) and duration distribution (B) for an individual subject and for different values of the threshold *θ* used to identify neuronal avalanches (Materials and Methods) (blue tick line: *θ* = 1 SD; green tick line: *θ* = 2 SD; red tick line = *θ* = 3 SD). The dotted black line is the power law fit for *θ* = 2*SD*. The average size as a function of the duration (C) follows the power law relationship 〈*s*〉 ∝ *T^k^* with *k* = 2 (black thick line) for all threshold values and for *T*’s smaller than the onset of the exponential cut-off of the duration distribution *P*(*T*). For larger T’s *k* = 1.3.

**Fig. S2:**
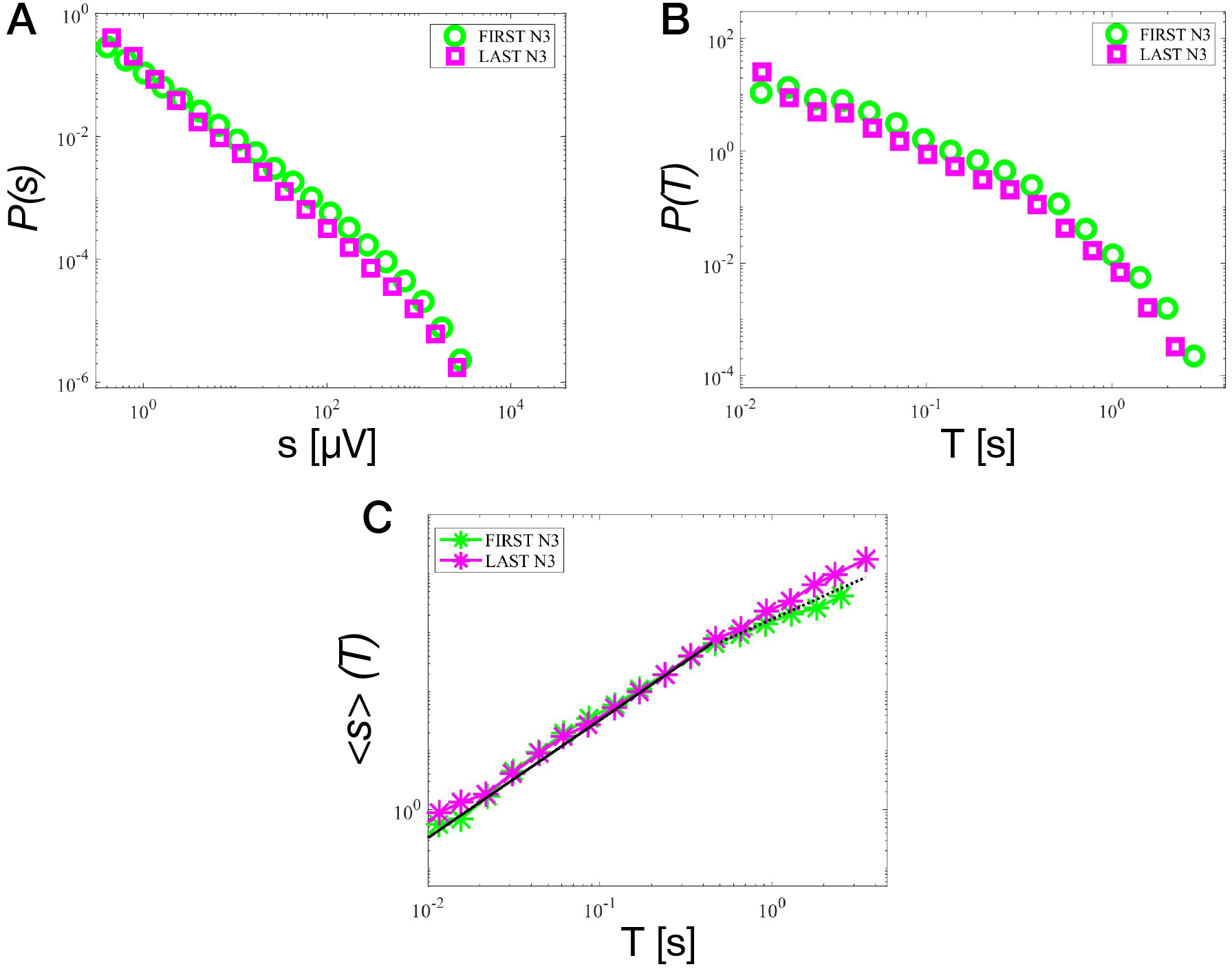
Distribution of size (A) and duration (B) for avalanches in the FIRST N3 (green) and the LAST N3 (magenta), and (C) average size as a function of the duration for avalanches in the FIRST N3 (green) and in the LAST N3 (magenta). Both the distributions and the relationship between average avalanche size and avalanche durations remain stable when moving from the FIRST N3 to the LAST N3.

